# UPhyloplot2: Visualizing Phylogenetic Trees from Single-Cell RNA-seq Data

**DOI:** 10.1101/2020.05.25.115550

**Authors:** Stefan Kurtenbach, Daniel A. Rodriguez, Michael A. Durante, J. William Harbour

**Affiliations:** Bascom Palmer Eye Institute, Sylvester Comprehensive Cancer Center, Interdisciplinary Stem Cell Institute, University of Miami Miller School of Medicine, Miami, FL, USA

## Abstract

Recent advances in single cell sequencing technologies allow for greater resolution in assessing tumor clonality using chromosome copy number variations (CNVs), which can be inferred from single cell RNA-seq (scRNA-seq) data using applications such as inferCNV. Inferences regarding tumor clonality are frequently visualized using phylogenetic plots, which previously required time-consuming and tedious manual analysis. Here, we present UPhyloplot2, a python script that generates phylogenetic plots directly from inferCNV output files. The tool is publicly available at https://github.com/harbourlab/UPhyloplot2/.

## Introduction

Single cell RNA sequencing (scRNA-seq) has become an important new tool for studying gene expression in individual cells of heterogenous samples. While this technology is still maturing, it is already providing powerful new insights into normal and diseased tissue types [1, 2]. In particular, single cell technology has resulted in great strides in cancer research. A hallmark of cancer is aneuploidy – an abnormal number of chromosomes or chromosomal segments – which often correlates with tumor aggressiveness [3–6]. Further, aneuploidy can be used to identify subclones of tumor cells and to infer tumor evolution, which can have important clinical implications [7]. Single cell sequencing can be used to analyze subclonal tumor architecture at unprecedented resolution [1, 8]. While single cell DNA sequencing (scDNA-seq) is an emerging technique for this type of analysis, it is very expensive and yet to be optimized. Alternatively, CNVs can be inferred from scRNA-seq using applications such as inferCNV [9], HoneyBadger [10], and CaSpER [11], using gene expression patterns to infer CNVs and to cluster cells into putative subclones. This approach for studying tumor clonality has been used successfully by our group and others [8, 12]. Tumor clonality is commonly used to visualize phylogenetic plots, where the length of tree branches is proportional to the number of cells in each subclone. Until now, such visualization required time-consuming and error-prone manual curation. Here we describe a new tool that we have created called UPhyloplot2, which is an enhanced version of UPhyloplot [13]. This application directly takes inferCNV output files and generates evolutionary phylogenetic plots.

## Results

UPhyloplot2 works directly with the output files of inferCNV, plotting evolutionary trees with the length of each branch correlating with the number of cells in the respective subclone. UPhyloplot2 was written in Python 3, making it easy to run on any platform (Figure 1). Uphyloplot2 uses the “HMM_CNV_predictions.*.cell_groupings” file generated by inferCNV (with HMM) to plot. Multiple “cell_groupings” files can be processed at once, to generate one output figure containing one tree per file. Subsequently, unique CNVs can be inferred using the “.HMM_CNV_predictions.*.pred_cnv_regions.dat” file. Four “cell_groupings” file being used as inputs (Figure 2). Output files are true SVG files, allowing for easy editing (colors, lines, branch rotation) in programs like Adobe Illustrator or any other SVG editor.

**Figure 1:**
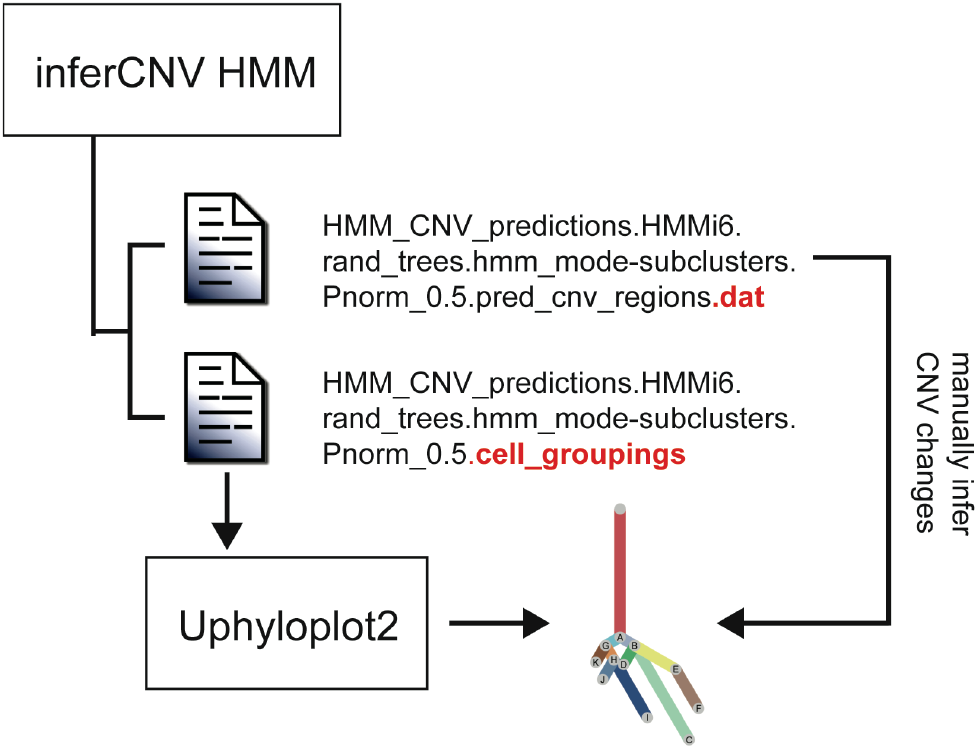
Generating phylogenetic tress with UPhyloplot2. The “cell_groupings” file is generated by inferCNV run with HMM, and used as inputs for UPhyloplot2. Unique CNVs for subclones can be inferred manually from the “.dat” file.

**Figure 2.**
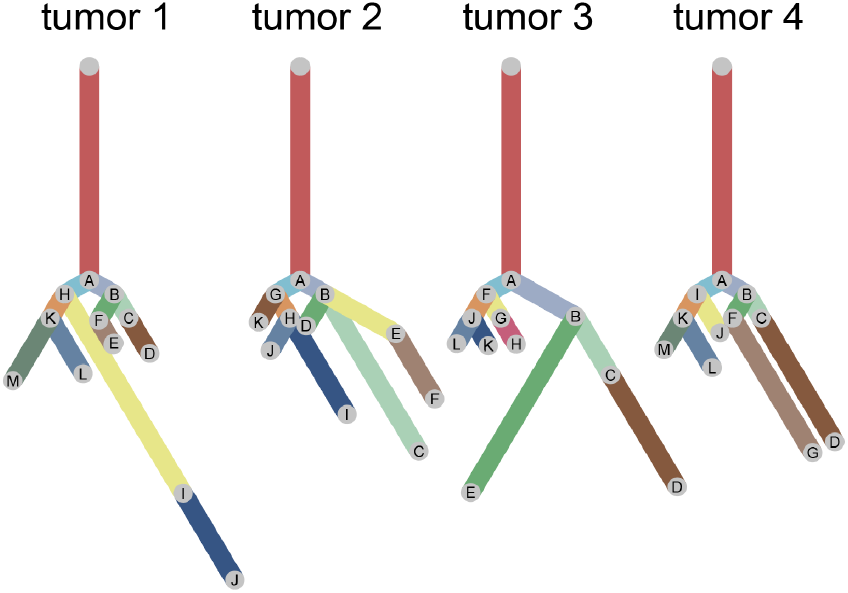
Example plots generated from four cell groupings files at once. Some branches were rotated to avoid overlapping.

## Discussion

The python script presented here allows phylogenetic trees of tumor subclones to be plotted from inferCNV output files. The script and further documentation are publicly available at https://github.com/harbourlab/UPhyloplot2/. In contrast to algorithms that estimate molecular time from whole-genome sequencing data using the number of mutations [14], the use of CNVs to infer clonality and tumor evolution is more complex because some chromosomal segments are selectively altered while others occur through massive genome reorganization such as chromothripsis [15, 16], chromoplexy [17] and anaphase catastrophe [18]. It is important to note that our methodology for plotting phylogenetic trees with branch lengths being proportional to the number of cells does not attempt to depict molecular time, but rather, the proportional size of each subclone. New methodologies are also being developed for analyzing single cell CNV and single cell mutation data [19]. In summary, we present an automated tool for generating phylogenetic trees from scRNA-seq data that allows the visualization of tumor subclones and heterogeneity.

